# HIV chromatin is a preferred target for drugs that bind in the DNA minor groove

**DOI:** 10.1101/618090

**Authors:** Donald W. Little, Samuel J. Schafer, John N. Anderson

**Affiliations:** Program in Biomedical Sciences, University of Michigan, dwlittle@umich; Department of Reproductive & Developmental Sciences, University of British Columbia; Department of Biological Sciences, Purdue University

**Author notes:** Contributed equally to this work with (Donald W. Little, Samuel J. Schafer).

**Keywords:** HIV, Nucleosome, Chromatin, Drugs, DNA minor groove

## Abstract

The HIV genome is rich in A but not G or U and deficient in C. This nucleotide bias controls HIV phenotype by determining the highly unusual composition of all majors HIV proteins. Since drugs that bind in the DNA minor groove disrupt nucleosomes on sequences that contain closely spaced oligo-A tracts which are prevalent in HIV DNA because of this bias, it was of interest to determine if these drugs exert this selective inhibitory effect on HIV chromatin. To test this possibility, nucleosomes were reconstituted onto five double-stranded DNA fragments from the HIV-1 pol gene in the presence and in the absence of several minor groove binding drugs (MGBDs). The results demonstrated that the MGBDs inhibited the assembly of nucleosomes onto all of the HIV-1 segments in a manner that was proportional to the A-bias, but had no detectable effect on the formation of nucleosomes on control cloned fragments or genomic DNA from chicken and human. Nucleosomes preassembled onto HIV DNA were also preferentially destabilized by the drugs as evidenced by enhanced nuclease accessibility in physiological ionic strength and by the preferential loss of the histone octamer in hyper-physiological salt solutions. The drugs also selectively disrupted HIV-containing nucleosomes in yeast as revealed by enhanced nuclease accessibility of the in vivo assembled HIV chromatin and reductions in superhelical densities of plasmid chromatin containing HIV sequences. A comparison of these results to the density of A-tracts in the HIV genome indicates that a large fraction of the nucleosomes that make up HIV chromatin should be preferred in vitro targets for the MGBDs. These results show that the MGBDs preferentially disrupt HIV-1 chromatin in vitro and in vivo and raise the possibility that non-toxic derivatives of certain MGBD might serve as a novel class of anti-HIV agents.

## INTRODUCTION

The tremendous genetic variation of HIV-1 is responsible for the appearance of drug and antibody resistant forms of HIV-1 that appear during infection and this genetic swarm is a major obstacle to the treatment of AIDS and to the development of an HIV vaccine. Viral genetic heterogeneity also produces variation in virulence, replication rates, cell tropisms and other properties (1,2). A drug designed to combat HIV should optimally target a viral feature that is conserved in this heterogeneous viral population, is critical for viral survival, acts on the integrated proviral genome, and is not found in the host cell. Although significant progress has been made during the past three decades in the development of anti-HIV drugs which has culminated in the use of combination antiviral drug therapy for treatment of viremia, HIV-drug-resistant mutants have been found for all nucleoside analogs and nonnucleoside and protease inhibitors used in the treatment of HIV-infected individuals. The emergence of multidrug-resistant variants of HIV-1 is also a concern for the treatment of the disease in the future (3,4).

The major challenge in current HIV research is the development of methods to eliminate the replication competent provirus that has integrated as a double stranded DNA molecule within the host cell chromatin. This viral reservoir is resistant to the host’s immune system and is resistant to the current drugs used to treat viremia. Upon session of antiviral drug therapy, competent proviral sequences are activated giving rise to new rounds of replication and the resumption of active infection. Little progress has been made during the past to eliminate or inactivate the replication competent provirus and there is no universal agreement as to the best strategies that should be used to approach the problem in the future (5,6).

The HIV genome and the genomes of other lentiviruses display an unusual variation in nucleotide composition being deficient in C but not G and rich in A but not U (7,8). We have previously shown that this nucleotide compositional bias is responsible the unusual composition of HIV proteins which rich in the polar amino acids encoded by A-rich codons and depleted in amino acids encoded by C-rich codons (8). The extreme amino acid compositions are characteristic of all major proteins of the viruses including the polypeptides that make up the hypervariable viral envelope and the conserved pol region that codes for reverse transcriptase and integrase. The global nature of these effects makes it likely that the variation in protein composition caused by the biased nucleotide frequencies is an important factor in determining the characteristic phenotype of HIV (8). The A-bias may also be a driving force that promotes enhance genetic variation of HIV and to increase the DNA curvature of the integrated viral genome that is caused by oligo A-tracts arranged in a 10 bp periodicity (8, 9). As a group, these observations suggest that the conserved A-bias of the HIV genome is critical for viral phenotype and thus the overall A-bias may represent a novel and fixed target for anti-HIV agents.

The minor groove binding drugs (MGBDs) are of interest in clinical medicine because of their antiviral, antimicrobial and antitumor activities (10-12). These ligands preferentially bind to AT-rich DNA sequences that are 4 or more bp in length. The biologically important targets of these drugs are poorly understood although many studies have focused on their abilities to inhibit the interaction of regulatory proteins to short oligonucleotide-length AT rich sequences. However, the importance of these targets in dictating the biologically relevant targets of these drugs is questionable because of the high frequency of isolated AT-rich oligonucleotide length sequences in the human genome. We previously reported a novel specificity of the MGBDs that requires much longer regions of DNA containing multiple closely spaced oligo A-tracts (13, 14). It was demonstrated that the drugs inhibited the assembly of such unusual DNA segments that are at least 100 bp long into nucleosomes. This inhibitory action may be relevant to HIV since the entire viral genome is A-rich and consequently nucleosomes containing HIV sequences may be targets for the MGBDs that are preferred over host sequences. The studies described in this report were designed to evaluate this possibility.

## EXPERIMENTAL PROCEDURES

DNA preparation and characterization—Plasmid PHRT25 (ref.15; from the American Tissue Culture Collection) containing the HIV-1 pol gene from isolate BH-5 was used as a template in the PCR to amplify HIV fragments 1-5. The nucleotide (NT) positions of the fragments in the HIV genome are 1973-2171, 2817-3024, 3458-3657, 4100-4303 and 4318-4519, respectively. The sequences of the top strand primers for fragments 1-5 were: 5’-CAATGGCCATTGACAGAAG, 5’-CAGAAAACAGAGAGATTC, 5’-GTCAATCAAATAATAGAGCAG, 5’-TTAAGACAGCAGTACAAATGGC, 5’-AGGTGAAGGGGCAGTAGT. The sequences of the bottom strand primers were: 5’-CCCAGAAGTCTTGAGT, 5’-TGCACTGCCTCTGTTA, 5’-CTAGCCATTGCTCTCCA,5’-TGCTGGTCCTTTCCAAAGTG, 5’-CTATAAAACCATCCCC. PCR amplification of the 2 Kb EcoRI pol fragment used the bottom strand primer for fragment 4 and the sequence of the top strand primer was: 5’-GGCCGGAATTCAATGGCCATTGACAGAAG. The amplified fragment was then cut at NTs 1973 and 4004 with EcoRI to give rise to the 2 Kb fragment which was inserted into the EcoRI site of plasmid P-2 as described below.

Two parent yeast shuttle vectors were used. P-1 is a pUC18 derivative containing yeast ARS 1 in the EcoRI site and the URA 3 gene in the HindIII site (16). P-2 is a pUC18 derivative with a 627 bp fragment from yeast ARS 2 in the Xho I site and the URA 3 gene in the HindIII site. HIV fragment 1 was blunt ended into the SmaI site of P-1 to give rise to P-1-H1 or inserted into the EcoRI site of P-2 to give P-2-H1. The 5 S rDNA sequence was inserted into the EcoRI site of P-2 to give P-2-5S rDNA. The 2Kb PCR amplified EcoRI fragment from HIV pol (1973-4004 bp) was inserted into the EcoRI site of P-2 to give P-2-H-2 Kb. To generate low copy number plasmids in yeast, a 289 bp Bam H1 fragment containing yeast CEN III was inserted into the Bam H1 site of each of the plasmids in the P-2 series.

In vitro studies-Synthetic (Syn # 61 and 67) and natural (5S rRNA gene from sea urchin) control 190-210 bp fragments were prepared as described previously (17). The synthetic sequences contain two regions of curved DNA of the form ((A5).(G/C5))4. Genomic DNAs from chicken erythrocytes and HeLa cells were digested with a mixture of HaeIII and Hinf I and fragments which comigrated with 200 bp standards were eluted from 1.4 % agarose gels and used for the studies in Figs. 3 and 6. DNA fragments were uniformly labeled with [alpha-32P] dATP or uniquely end-labeled by using one 32P-end-labeled primer in the PCR. Genomic DNAs were end-labeled by standard procedures. DNA fragments were purified by electrophoresis through agarose gels and appropriate bands eluted from gels by the crush-soak method.

MGBDs were from Sigma and drug solutions in 10 mM Tris-HCl (pH 7.5) containing 10 mM NaCl were stored in the dark at —70°C. Nucleosomes were reconstituted onto the DNA fragments by the standard exchange-salt dilution method (1.0 M NaCl to 0.1 M NaCl) using H1/H5 deficient nucleosomes from chicken erythrocytes as a source of core histones (17). For translational mapping studies with HIV fragment 1, the reconstituted nucleosome was digested with exonuclease III and analyzed as detailed previously (17). The major ExoIII pause sites were found at 2017 and 2161 bp with a midpoint of 2089 bp. The predicted position of the dyad from a previous computational analysis of DNA curvature was 2092 bp (9). For footprinting analysis of HIV fragment 1, DNase I digestion of the DAPI-DNA complex was carried out for 3 min using 8 U/ml as described previously (13). For the restriction enzyme studies in Fig. 6, digestions were carried out for 40 min at 37°C in 10 mM MgCl2 containing 40 mM NaCl and the fragments analyzed on 8% native PA gels. The nitrocellulose filter binding assay was carried out as described previously (18) using the NaCl concentrations for sample application and washing that were the same as those used in Fig. 5. The relative binding affinities of DAPI for HIV and control DNAs were determined by fluorescence-enhancement at 465 nM as detailed previously (13).

Studies using yeast-S. cerevisiae strain J17 (MAT a,his2,ade1,trip1,met14,ura3-52) was used as the host for plasmid transformation. The isogenic derivative containing the rad 52 deletion was obtained from K.S. Bloom (19). Plasmid DNAs were used to transform yeast to uracil prototrophy and transformed cells were grown at 37°C in synthetic medium lacking uracil (16). For chromatin studies, cultures (100-200 ml) were diluted to 0.2 A 660/ml, drugs were added as indicated and the cells grown overnight to late log phase corresponding to an A660/ml of 1.3-2.0. DAPI (4 uM) reduced growth rates by approximately 20% while concentrations > 10 uM caused severe growth retardation. Berenil at concentrations up to 150 uM was without effect on the generation time. Spheroplasts were isolated essentially as described by Kent et al. (20) except Lyticase (Sigma) was used in place of Glusulase and spermidine was omitted from the MNase digestion buffer. Spheroplasts were permeabilized with NP-40 and digested with MNase at 37°C. for 6 minutes. For indirect-end labeling, DNA obtained from MNase digested spheroplasts was secondarily digested with Ava II and fragments were electrophoresed on 1.0 % agarose gels, transferred to Zeta probe membranes (Bio-Rad) in 0.4 M NaOH and the membranes hybridized with multiprime labeled 200-300 bp probes that abutted the Ava II restriction sites. Gels were calibrated using plasmid restriction fragments which are not shown in the figures. For the experiments in the top panels of Fig. 7, blots were probed with HIV sequences or with a 190 bp Msp1 fragment from pUC18 which is located 734 bp from the HIV DNA. Blots from chloroquine gels were probed with pUC18. For analysis of DNA topoisomers, DNA was prepared from 10 ml cultures as described previously (16) and electrophoresed on 0.9-1.1 % agarose gels in Tris-phosphate-EDTA in the presence of 1.5 ug/ml of chloroquine (21). Under these conditions, the bulk of the topoisomers are negatively supercoiled. Values for drug-induced loss of superhelical turns similar to those given in Fig. 8 were obtained when 90 ug/ml chloroquine was used. Under these conditions, the bulk of the topoisomers were positively supercoiled, reflecting the differences in negative supercoiling (22). Plasmid copy number was determined as described previously (16).

## RESULTS

There are several characteristics of the HIV-1 genome which should make it a preferred target for the MGBDs. First, as shown in Fig. 1, approx. one fourth of the nucleotides in the genome are AT bps in sites that are at least four bp in length: the minimal binding-site size for most of the MGBDs. The coding strands of these AT sites are A-rich (mean~70% A) reflecting the compositional bias of the genome. The level of AT sites (Fig. 1A) also follows the A-bias along the genome (7,8) being high in gag, pol and env regions and reduced only in regions of overlapping reading frames and in the long terminal repeats. There are multiple nucleosome-length stretches in pol and env with more than 40% of their AT bps in AT sites. On average, there is one ~4 bp AT sequence per 12 bp of DNA in these regions. Nucleosomes that contain such high levels of AT sites should be drug sensitive provided that the AT sequences have narrow minor grooves (13,14). The virus does not select against these high AT site densities since the levels of AT sites in the genome and in these subgenomic segments are about 1.2-1.3 fold greater than those in randomized HIV sequences. Taken together, these results imply that the A-bias unopposed by selection forces favors a high frequency of potential drug binding sites in the HIV genome.

**Fig. 1.**
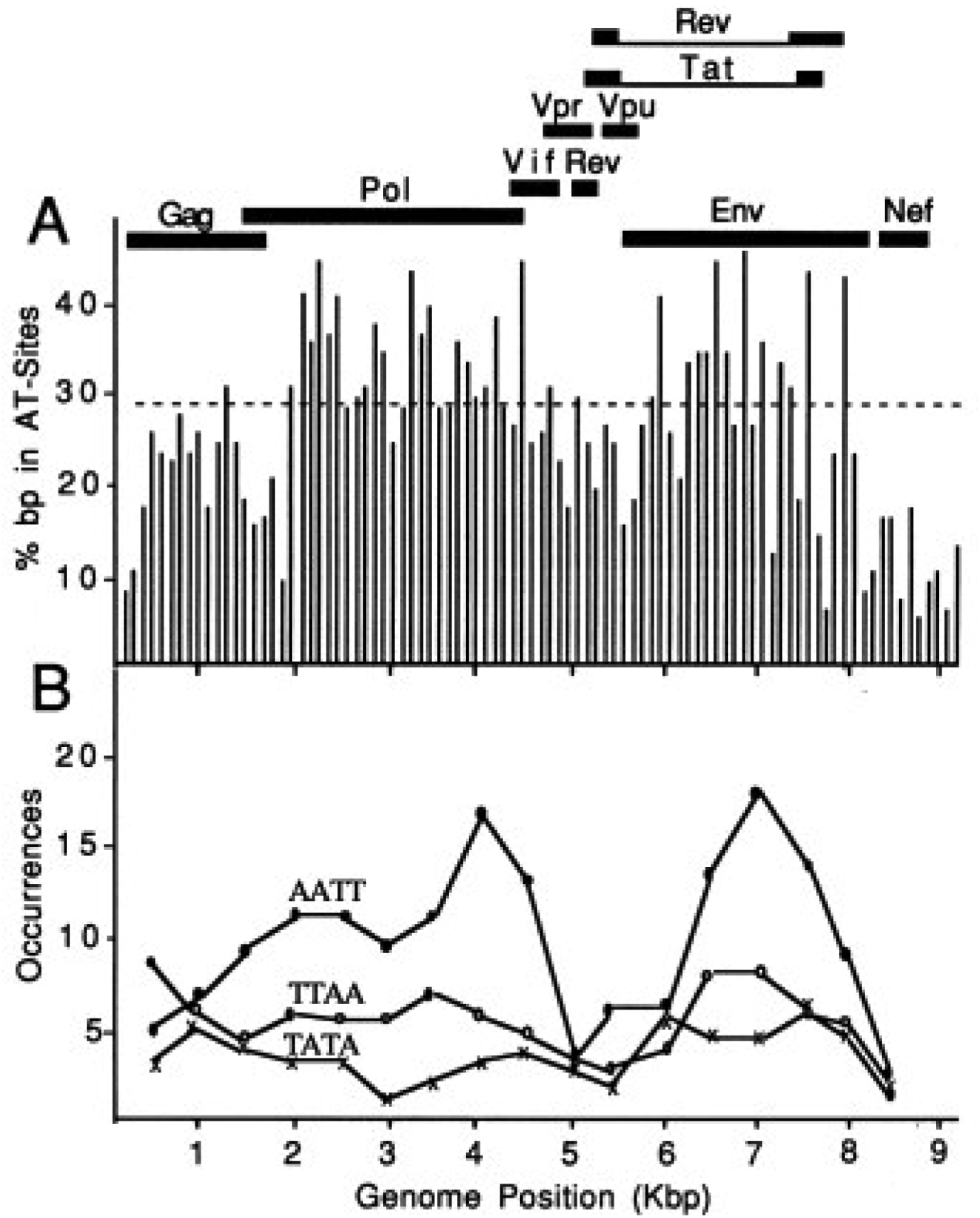
AT-tract density in HIV DNA. Panel A: The percentage of nucleotides in 200 bp segments (window step = 100 bp) along the HIV genome that are in AT sites of 4 or more bp in length are plotted in the diagram. The dotted line indicates the 29% AT-site density displayed by HIV fragment 5 (Table 1). The corresponding values for the control 5S rDNA sequence, Syn 61 and Syn 67 (Figs. 3 & 5) are 17 %, 20 % and 20 %, respectively. Panel B: Occurrences of AATT (solid circles), TTAA (open circles) and TATA (x) in the coding strand were determined at a window width of 1000 nucleotides and window step of 500 nucleotides.

**Table 1.** Characteristics of the AT and GC Sites ≥ 4 bp in five HIV *pol* fragments.

|  | Fragment |  |  | AT-Sites |  |  |  | GC-Sites |  |  |
| --- | --- | --- | --- | --- | --- | --- | --- | --- | --- | --- |
|  | bp | Position* | %A+T | N | bp | %bp | %A** | N | bp | %bp |
| 1. | 199 | (1973-2171) | 67 | 16 | 5.5 | 44 | 69 | 2 | 4.5 | 4 |
| 2. | 206 | (2817-3024) | 62 | 14 | 5.4 | 37 | 71 | 3 | 4.3 | 6 |
| 3. | 198 | (3458-3657) | 63 | 12 | 5.8 | 35 | 67 | 1 | 4.0 | 2 |
| 4. | 202 | (4100-4303) | 64 | 15 | 5.1 | 38 | 66 | 4 | 4.5 | 9 |
| 5. | 200 | (4318-4519) | 62 | 12 | 4.8 | 29 | 63 | 2 | 4.0 | 4 |

\* Position in isolate BH-5
\*\*The %A is given as coding strand composition.

Fig. 2 shows the composition of all AT tetranucleotide sequences in the HIV-1 genome (A), the HIV pol region (B), and the five pol fragments (C) used in the studies described below. The effect of the genomic bias on the composition is clearly evident with the level of homopolymeric tracts of A being about 3-fold and 15-fold greater than those of T and C, respectively. The underlined sequences lack a TA step and these sequences are the most prevalent in the mixed AT sequences at each frequency of A in the three sequence sets. Each of these sequences is also overrepresented in HIV DNA relative to randomized HIV DNAs as shown by the numbers in panels A and B which represent the ratios of the occurrences in the natural to the randomized sequences. These findings may be significant since the TA step widens the minor groove of AT sites (23-26) and the compressed groove is required for high affinity drug binding and nucleosome disruption by the MGBDs (10, 13, 27, 28). In contrast, most of the sequences with TA dinucleotides are underrepresented in HIV DNA relative to the shuffled sequences and this negative selection is particularly pronounced in the tetranucleotide with two TA steps and those with a single TA step in the central region. The relative occurrences of the sequences that make up the AT sites with 50 % A were AATT >> ATTA > TTAA ~ TAAT > ATAT ~ TATA. This order is inversely related to the DNA minor groove width of these sequences (23-26) and parallels MGBD binding affinity (13, 27, 28) and the effectiveness of the drugs for disrupting nucleosomes on DNA that is rich in AT-sites (13,14). The overrepresentation of AAAA and AATT and other sequences that lack TA dinucleotides also follow the occurrences of the AT sites along the viral genome while the sequences with TA steps display occurrence patterns that are lower and more uniform throughout the viral genome (Fig. 1B). Consequently, the segments containing clustered AT sites in pol and env of HIV should be especially sensitive to the MGBD.

**Fig. 2.**
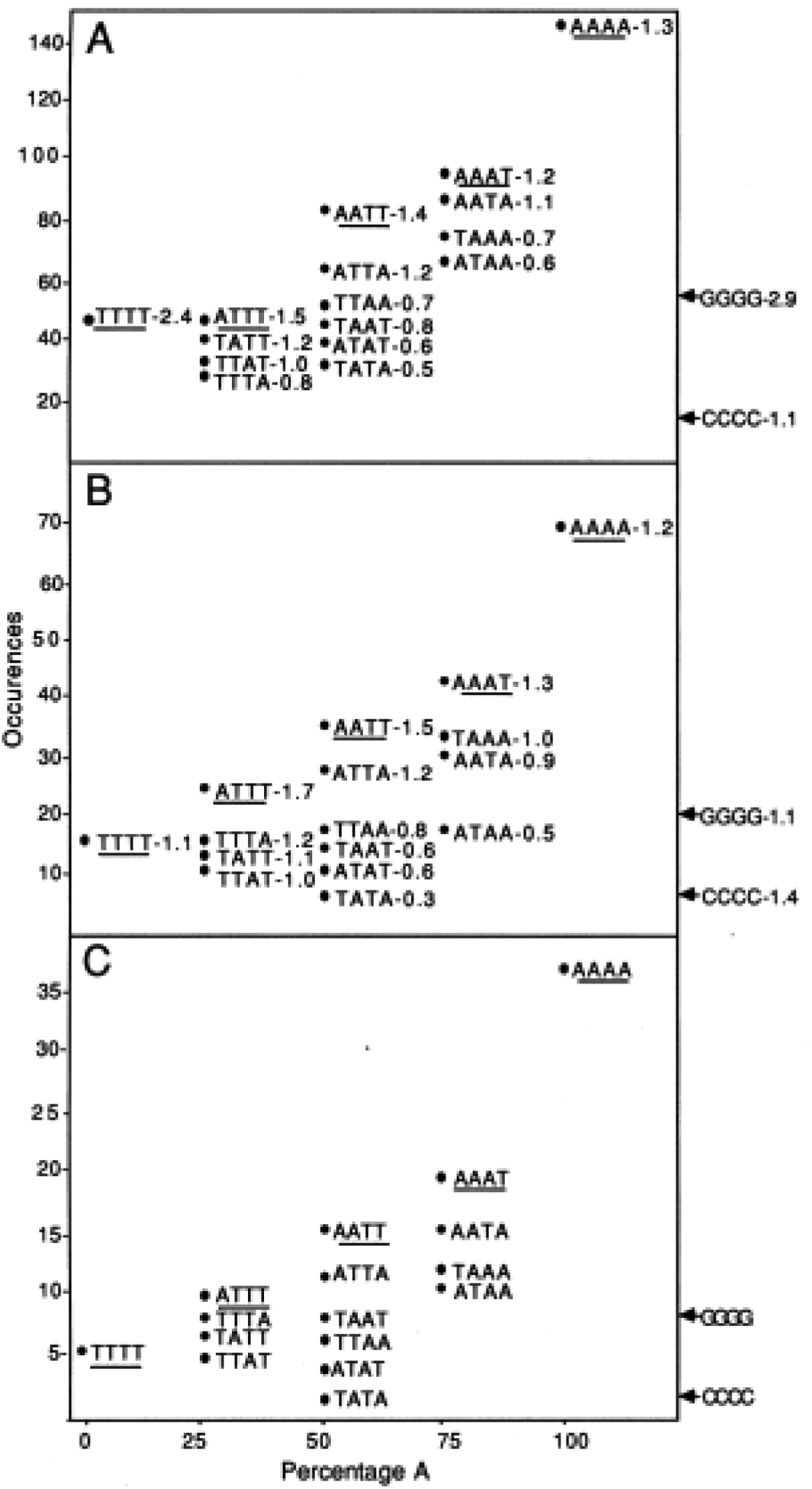
AT tetranucleotide sequences in HIV-1 Occurrences of all AT tetranucleotide sequences in the coding strand of the HIV-1 DNA genome (A), the HIV pol region (B), and the five pol fragments (C) described in Table 1 are plotted against the percentage of A in each region. The numbers adjacent to the tetranucleotide sequences in A and B are the ratios of the occurrences in the natural to the randomized HIV sequences. The underlined sequences lack a TA step.

Table 1 gives characteristics of the five DNA fragments from HIV pol that were used in the studies described below. The fragments extend from the beginning of reverse transcriptase coding region into the viral integrase gene. The ~200 bp fragments have a high density of AT-sites > 4 bp in length. On average, each has 14 AT sites that are 5.3 bp in length with 37% of nucleotides residing within these sites. These observed frequencies of site numbers and densities are approximately 1.3 fold higher than those in the randomized sequences. The frequency of the AT-sites is about 7-fold greater than the frequency of GC sites that are > 4 bp in length and these GC bps make up only 5 % of the nucleotides in the sequences. Each of the fragments also shows enrichment of AT sites that are not interrupted by TA steps similar to the sum of the sequence set which is shown in Fig. 2C.

Since MGBDs inhibit the assembly of nucleosomes on sequences with multiple closely spaced oligo-AT sequences with narrow minor grooves which serve as drug binding sites (13, 14), it was of interest to determine if the drugs exert their selective inhibitory action on HIV chromatin. To study this possibility, nucleosomes were reconstituted onto the five fragments in the presence and absence of several MGBDs. Free DNA was then separated from nucleosome DNA by electrophoresis on polyacrylamide-glycerol gels. A sample of the data is shown in Fig. 3 and the results of several similar experiments are summarized in Fig. 4. The studies demonstrated that the MGBDs inhibited the assembly of nucleosomes onto all five of the HIV segments. The marked specificity of the drugs for HIV DNA is illustrated by the results in Fig. 3 which show that these ligands had no detectable effects on the assembly of nucleosomes onto a control fragment and genomic DNA. For all HIV fragments, the relative order of inhibition was DAPI (4,6-diamidino-2-phenylindole) > distamycin > Hoechst 22358 > berenil >> pentamidine. For all drugs, the relative order of inhibition was HIV fragment 1 > 2, 3, 4 > 5. This order parallels the density of AT sites > 4 bp as can be seen from the data in Table 1. This finding is in agreement with our previous studies which demonstrated that drug sensitivity is directly related to the frequency of AT sites in nucleosome-length DNA (13, 14). Selective inhibition of nucleosome formation was seen with HIV fragment 5 which contains 29% of nucleotides in AT sites. As indicated by the dotted line in Fig. 1A, approximately 65% of all of the possible 200 bp segments in HIV pol and 35% of all of the possible 200 bp segments in the entire viral genome have AT site densities that are equal to or greater than this level. These same sites also tend to be enriched in AT sequences without TA steps as shown by the example given in Fig. 1B. These observations suggest that a relatively large fraction of the nucleosomes along HIV chromatin should be a preferred target for MGBDs.

**Fig. 3.**
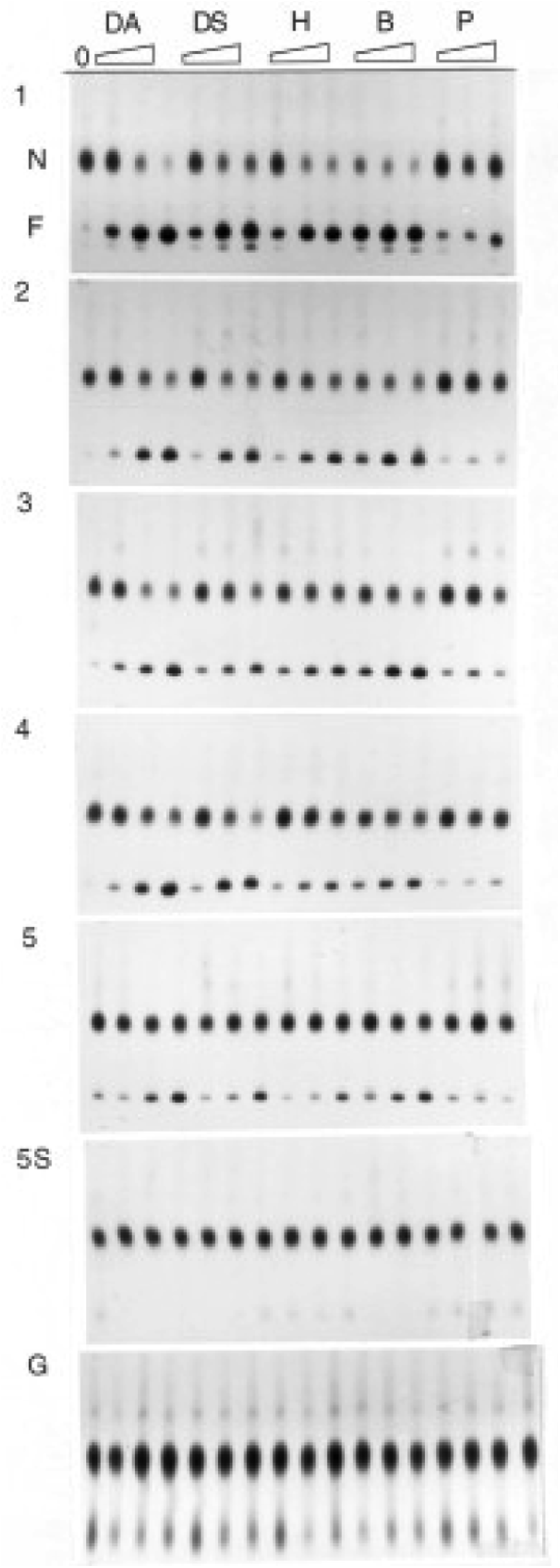
Selective inhibition of nucleosome assembly on HIV pol fragments by MGBDs. Labeled HIV fragments 1-5, the 5 S rDNA sequence and genomic (G) DNA from HeLa cells were incubated in 1 M NaCl with increasing amounts of MGBDs (1, 5 and 10 uM for DAPI (DA), Hoechst (H) and distamycin (DS); 2,10 and 20 uM for berenil (B) and pentamidine (P)) prior to addition of 300 ng of erythrocyte nucleosomes. The salt concentration was reduced by stepwise dilution and the products were electrophoresed on PA-glycerol gels in order to resolve nucleosomal (N) and free (F) DNA.

**Fig. 4.**
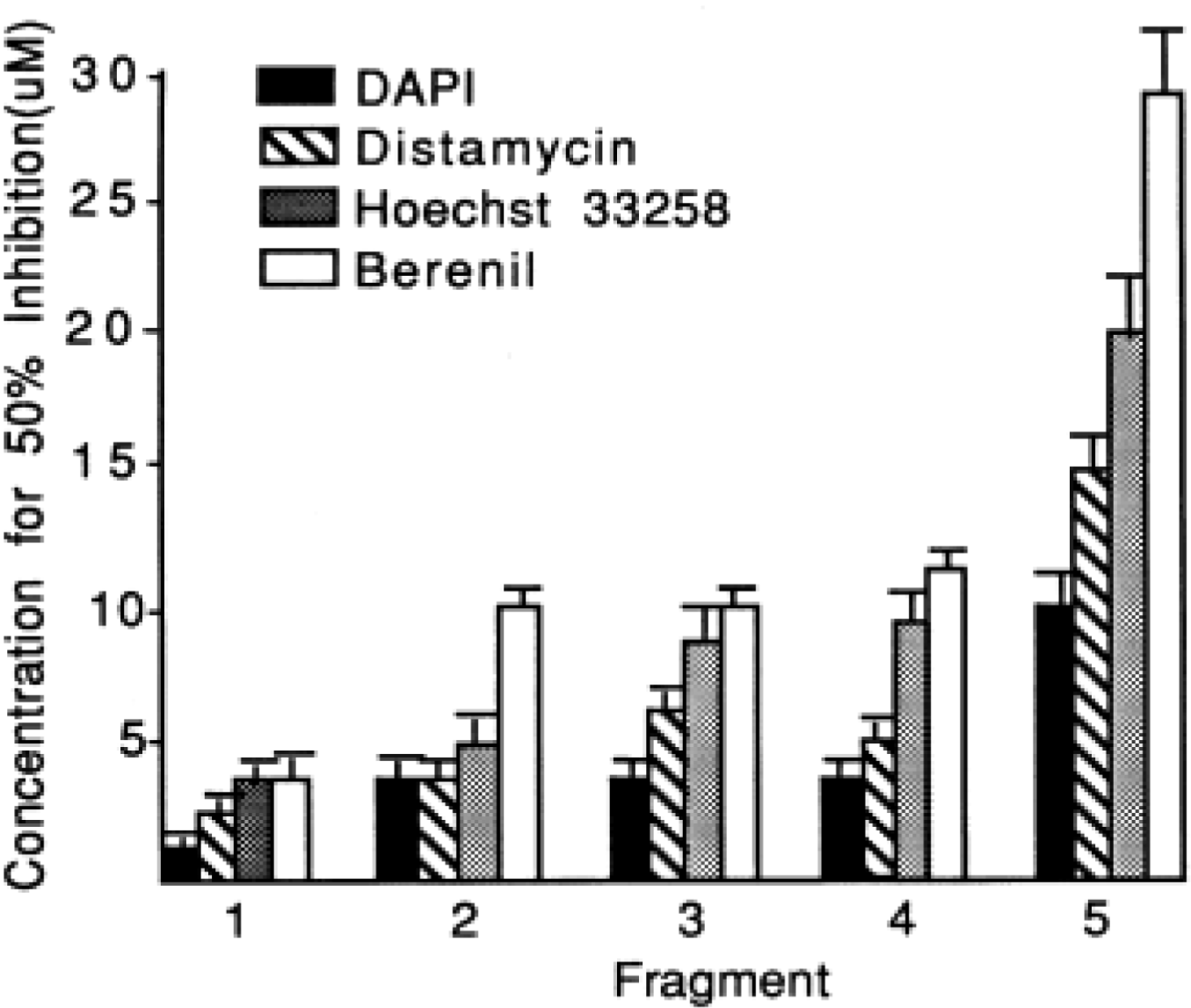
Effectiveness of the MGBD on inhibiting nucleosome formation on the five HIV 1 fragments. Mean (+ S.E.M.) micromolar concentrations of drugs that caused a 50 % inhibition of nucleosomes formation was determined from densitometric scans of the results of 4-6 experiments of the type shown in Fig. 3.

Previous studies have shown that the MGBDs promoted the destabilization of nucleosomes that were pre-assembled onto DNA molecules rich in AT sites and this effect was most pronounced at high ionic strength (13,14). This approach was taken in the present study to determine if MGBDs destabilize nucleosomes assembled onto the five HIV fragments. Fig. 5 shows the fate of preassembled nucleosomes containing the five HIV sequences and several control DNAs when they were incubated for 1 hour at 37° C in 0.1-0.5 M NaCl in the presence and in the absence of DAPI. Nucleosomes assembled onto the HIV DNAs were preferentially destabilized by the drug and the effect was most apparent in 0.5 M NaCl where up to 95% of the histone octamers were lost from DNA. The relative order of drug-induced nucleosome loss (fragment 1 > 2, 3, 4 > 5) was the same as the order for drug-induced inhibition of nucleosome assembly and density of AT sites as can be seen by comparing the results in Table 1 and Figs. 4 and 5. The template selectivity is also clearly evident from the results since octamers were retained on all control fragments and genomic DNA in all salt levels.

HIV fragment 1 contains two strongly curved DNA segments of ~50 bp separated by a ~30 bp region of low curvature (9). This pattern of curvature is characteristic of nucleosome positioning sequences (17, 29). The central position of this nucleosome along with the positions of restriction sites for HaeIII and RsaI is shown in the map at the bottom of Fig. 6. In order to see if a MGBD destabilizes this nucleosome in low ionic strength, a preassembled nucleosome containing this sequence was incubated for 40 min at **37°** C in 40 mM NaCl in the presence and in the absence of DAPI. Under these conditions, no more than 5% of the DNA was rendered histone-free by drug treatment. Restriction enzymes were then added and the reaction terminated after an additional 40 min. In the absence of the drug, HaeIII site 1 which lies outside of the major positioned core was cleaved extensively while HaeIII site 2 and the RsaI sites were more protected from enzyme cleavage. The analysis demonstrated that DAPI increased the extent of cleavage of these internal nucleosome sites by 3-8 fold (Fig.6).

**Fig. 5.**
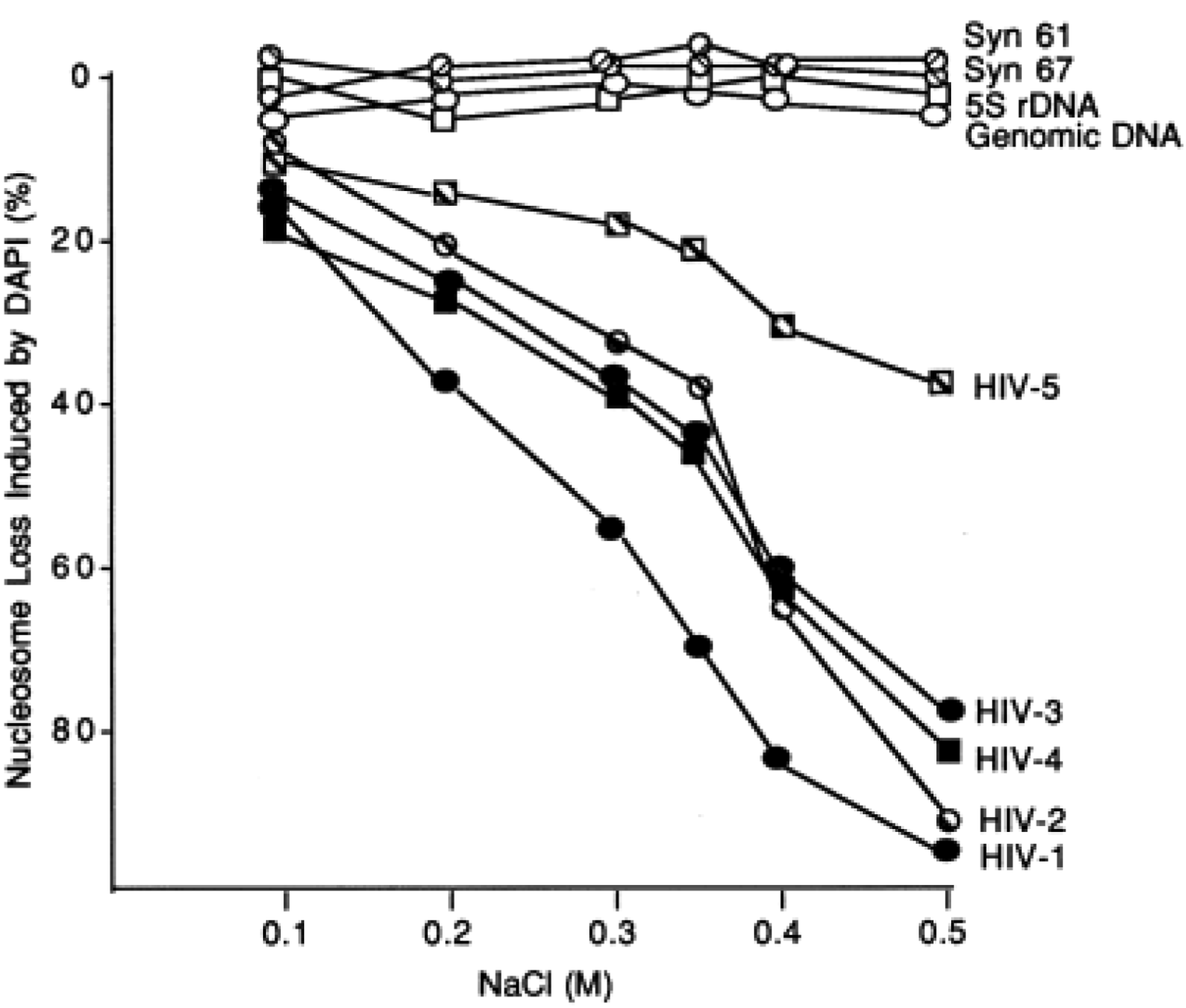
Selective destabilization by DAPI of nucleosome preassembled on HIV fragments. Labeled HIV fragments 1-5, the 5 S rDNA sequence, synthetic DNA fragments (Syn) 61 and 67 and genomic DNA from chicken were assembled into nucleosomes as in Fig. 3 and then incubated in the absence and in the presence of 10 uM DAPI at the indicated NaCl concentrations for 1 hour at 37°C. The percentage of the nucleosomes that were lost in the presence of the drug was then determined following electrophoretic analysis. With all sequences, more than 90% of the nucleosomes were retained in the absence of drug in all salt levels.

**Fig. 6.**
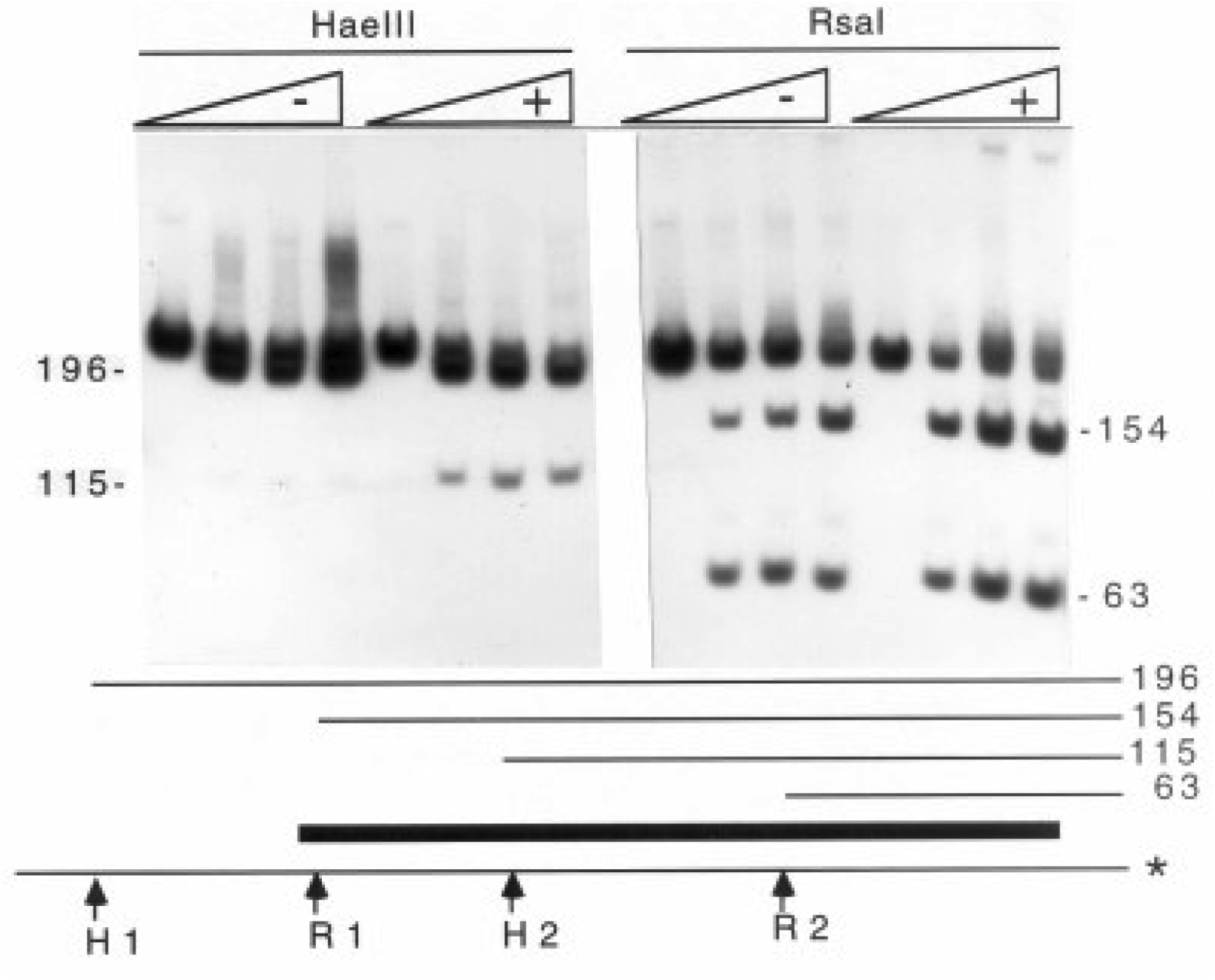
Effects of DAPI on the restriction enzyme accessibility in reconstituted nucleosomes. A schematic diagram of the major position of the single nucleosome (dark bar) and the restriction sites for HaeIII (H 1 & 2) and RsaI (R 1 & 2) in HIV fragment 1 is shown at the bottom of the figure. In the gel, a preassembled nucleosome containing this sequence labeled on the bottom strand (*) was incubated for 40 min at 37°C in 40 mM NaCl and 10 mMMgCl2 in the absence (-) and in the presence (+) of 10 uM DAPI. HaeIII (left 8 lanes; 0, 50, 150, 375 U/ml) and RsaI (right 8 lanes; 0, 50, 150, 375 U/ml) were then added and the reaction terminated after an additional 40 min.

The above studies established the selective inhibitory effects of the MGBDs on HIV-1 containing nucleosomes in vitro. To provide a first step to evaluate the effects of the drugs on HIV chromatin in vivo, HIV segments were cloned into yeast shuttle vectors and cells transformed with the plasmids were grown for about 20 hours in the absence and in the presence of 4 uM DAPI. Permeabilized spheroplasts were digested with micrococcal nuclease (MNase) and the digested DNA was electrophoresed, blotted and probed with HIV DNAs. The MNase ladder observed in the absence of drug is indicative of regularly spaced nucleosomes and the definition of the nucleosome repeat was reduced by DAPI as seen in Fig. 7. Reduction in the resolution of the ladders was seen with the chromatin assembled onto the 199bp HIV segment 1 in two different sequence contexts (panels A and B). Partial but not complete loss was noted in the chromatin assembled onto the longer 2Kb HIV segment and this reduction was more clearly seen in the minimally digested samples (panel C). This result is expected since some but not all of the nucleosomes assembled on this long sequence are expected to be drug sensitive. The drug had little effect on the nucleosome ladders containing bulk yeast DNA and little or no effect on the nucleosome profile of sequences from the blot probed with a segment of pUC18 (panels D, E). The reduction in the nucleosomal repeat in panels A-C is interpreted as representative of either the preferential loss or disruption of nucleosomes along the HIV sequences.

**Fig 7.**
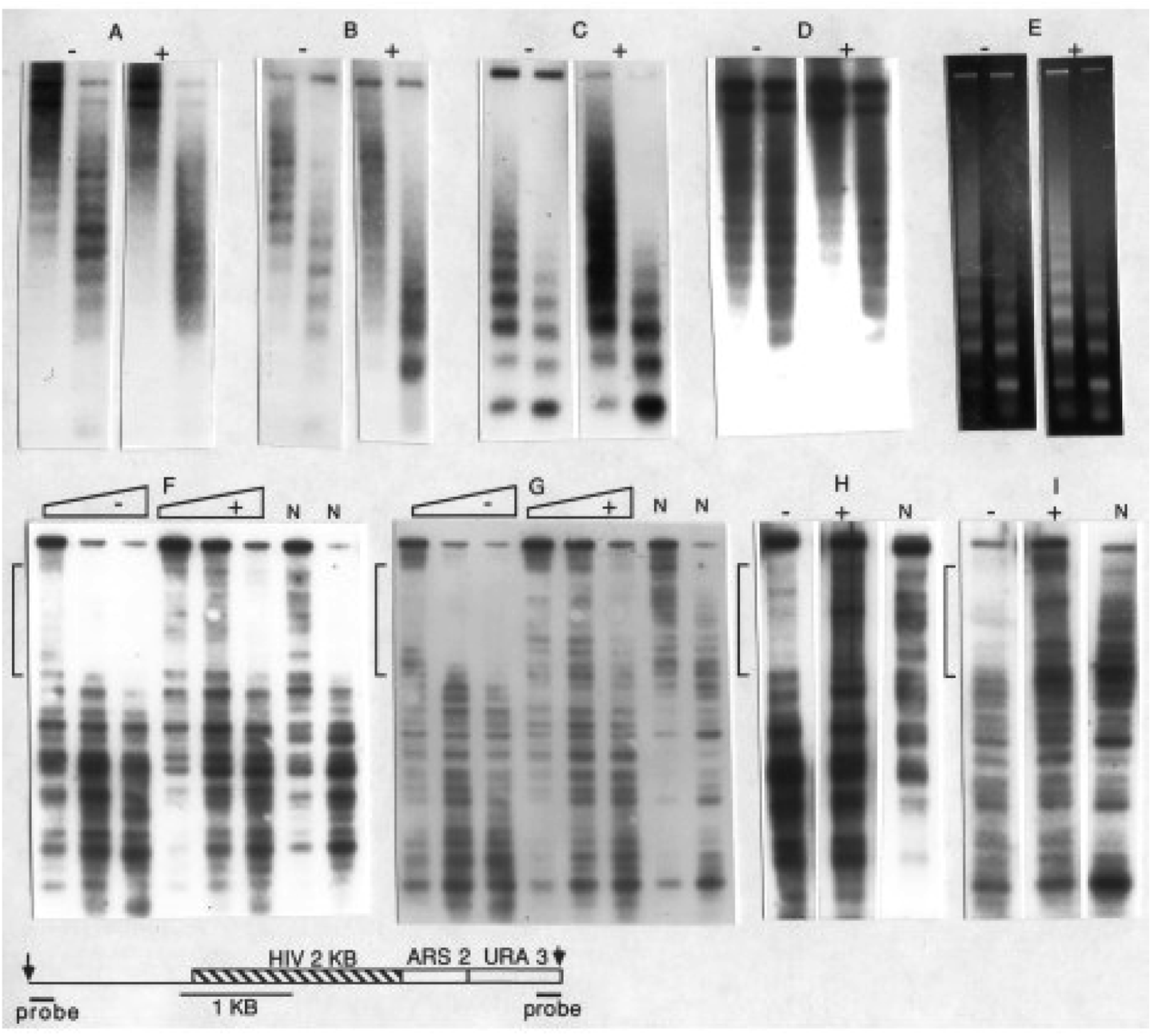
Top: Cells carrying plasmids with HIV sequences (A-D, P-1-H1, P-2-H1, P-2-H-2Kb, P-2-H-2kb, respectively) were grown for about 20 hours in the absence (-) and in the presence (+) of 4 uM DAPI. Permeabilized spheroplasts were digested with MNase (100 and 350 Units/ml) and the digested DNA was electrophoresed, blotted and probed with the HIV DNA in the corresponding plasmids (A-C) or puc18 DNA (D). An ethidium bromide stained gel from cells with P-2-H2Kb is shown in (E). Bottom: Indirect end-labeling analysis was performed on MNase digests obtained from cells that carried the longer HIV segment. The map of the region indicates the positions of the AvaII sites (arrows), hybridization probes, pUC18 DNA (single line), the 2 Kb HIV segment and a portion of the URA3 gene. The sequence that corresponds to the 199bp HIV segment 1 is on the left of the 2 Kb segment. Chromatin samples from control (-) and DAPI treated (+) cells were digested with increasing amounts of MNase (50, 100 and 200 U/ml). DNA was then digested wi th AvaII, electrophoresed on 1% gels and blots probed with the URA 3 probe (F) and the pUC18 probe (G). Chromatin samples from control (-) and berenil treated (+) cells were digested with MNase (100 U/ml) and the blots probed with the URA 3 probe (H) and the pUC18 probe (I). Naked DNAs carried through the procedure are indicated by N. The marked of MNase on naked AT-rich DNA seen on these lanes is well known The approximate gel positions of the 2 Kb HIV segment are indicated by the vertical brackets.

To further characterize this effect, indirect end-labeling analysis was performed on MNase digests obtained from control and DAPI treated cells that carried the longer HIV segment (Fig. 7, F, G). In the absence of drug, the HIV sequences resided within a nuclease protected region which is nuclease sensitive in naked DNA. DAPI enhanced the nuclease cutting of the HIV sequences in chromatin rendering the digestion pattern similar, but not identical, to that seen in the protein-free samples. The cutting pattern of sequences adjacent to the HIV segment were similar in the presence and in the absence of the DAPI providing evidence for the template selectivity of this drug action. Essentially the same results were seen in experiments where 60uM berenil was used in place of DAPI (Fig. 7. H, I).

In order to provide an alternative strategy to study the effects of the MGBDs on the in vivo assembled chromatin, DNA topoisomers from drug-treated and control cells were resolved in agarose gels containing chloroquine (Fig. 8). DAPI promoted a topological change toward a loss of superhelical turns of the HIV-containing plasmids while having little or no effect on control plasmids that lacked HIV sequences. Such a reduction is most often interpreted as a loss of nucleosomes since each nucleosome induces a single negative superhelical turn in a closed circular DNA molecule (21, 32). Densitometric scans of these and other autoradiograms indicated an average loss of 1-2 nucleosomes on the plasmids with the ~200 bp HIV sequence 1 and up to 4-6 nucleosomes on the plasmid containing the larger segment of HIV DNA as seen in the histogram in Fig. 8. The value for the larger segment is significantly less than the 13 nucleosomes that should be present on this sequence in the absence of drug which implies that a fraction of the nucleosomes on this HIV segment are drug resistant. This result is consistent with the partial loss of the nucleosome ladder induced by drug treatment (Fig. 7C) and with calculations of the AT-tract densities of 200 bp segments that make up this fragment which indicate that only about 2/3 of the nucleosomes in this region of HIV should display detectable drug sensitivity (Fig. 1A). The results shown in Fig. 8 (bottom panel) also demonstrate that DAPI was several fold more effective than berenil at inducing nucleosome loss in vivo in agreement with the relative potencies of these drugs determined by the in vitro studies shown in Figs. 4 and 5.

**Fig. 8.**
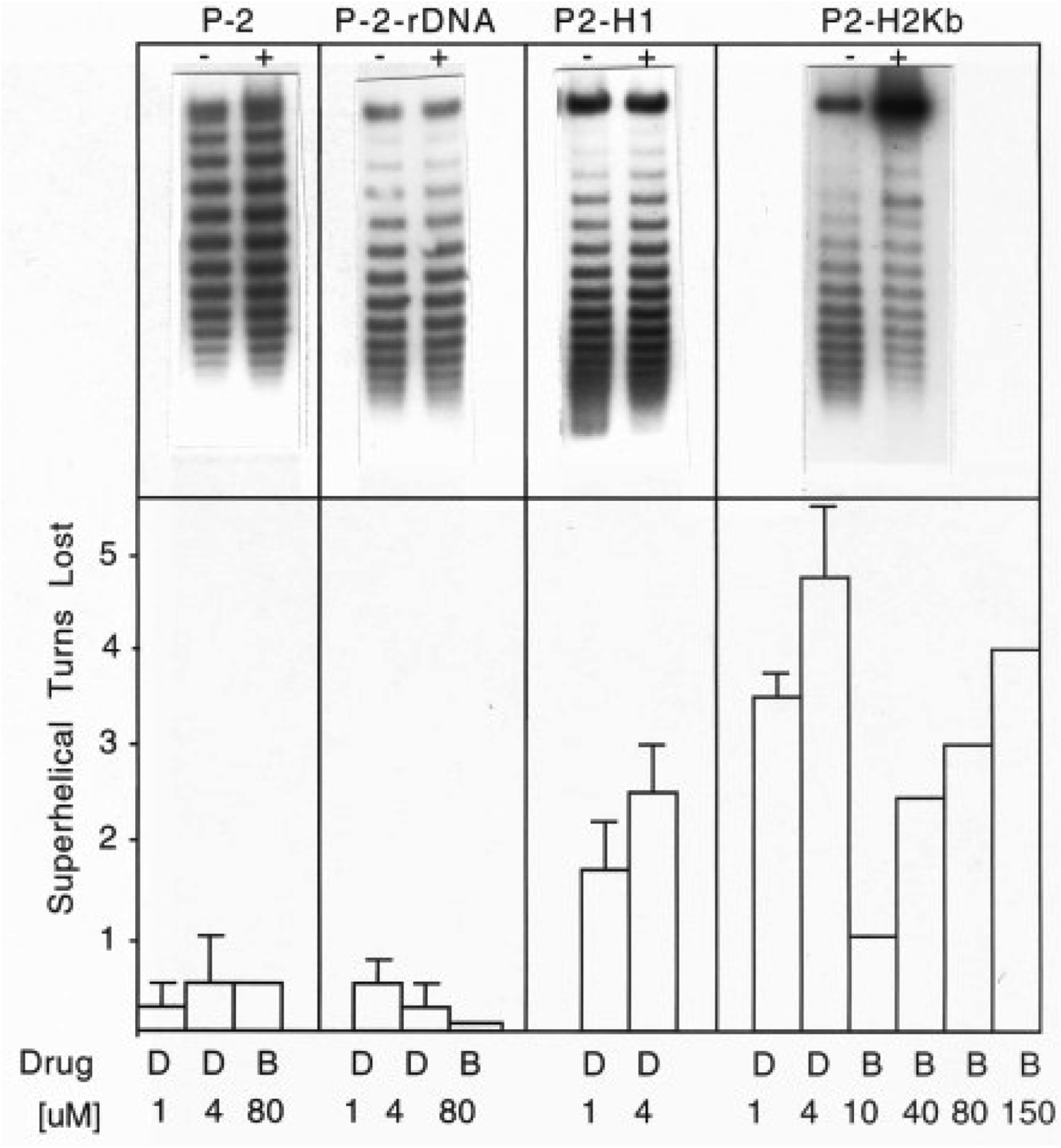
Effects of DAPI and berenil on the superhelicity of plasmids in yeast. Top: Cells that carried the indicated plasmids were grown in the presence (+) and in the absence (-) of 4 uM DAPI for 18 hours and the distributions of topoisomers were analyzed in chloroquine-agarose gels as described in Experimental Procedures. The DNA at the top of the gels, which is particularly prevalent in the P2-H2kb +DAPI lane, represent relaxed/nick circular plasmid (21, 32). Bottom: Mean peak ladder positions (+/-S.E.M.) obtained from densitometric tracings of 2-6 independent experiments of the type shown in the top panels were used to calculate the loss of superhelical turns induced by drug. Error bars are not given for the experiments performed only twice. The drug (D, DAPI; B, berenil) concentrations in uM are indicated

Since DNA that is devoid of nucleosome is hypersensitive to nucleases (32,33), it was of interest to determine if loss or disruption induced by a MGBD might render the HIV DNA nuclease hypersensitive in vivo. To test this possibility, HIV fragment 1 and the longer HIV segment on centromere-containing plasmids were introduced into yeast cells that lacked the RAD 52 gene, which is required for repair of DNA double-strand breaks (34). DAPI and berenil had no effect on plasmid copy number or HIV sequence integrity in these cells or in wild-type cells nor did the drugs preferentially affect the growth rate of these cells in selective media lacking uracil relative to those that were transformed with control plasmids that lacked the HIV DNA. Likewise, the drugs did not increase the ratio of relaxed circular/supercoiled DNA of the HIV-containing plasmids. Thus, it appears that nucleosome loss in vivo does not necessarily render the HIV DNA sensitive to cleavage by cellular nucleases under the conditions used in these studies.

## DICUSSION

Most of the parent MGBDs were developed as trypanocides and berenil remains used for this purpose today in Africa and Asia. Pentamidine has also been used to prevent and treat opportunistic pneumonia that is caused by Pneumocystis carinii which is common in AIDS patients.. The drugs have found additional applications in clinical medicine including use as general antitumor, antibacterial and antiviral agents (12, 35-38). However, the targets and modes of action of the drugs remain, for the most part, unknown. Although the MGBD distamycin and related analogues show anti-HIV activity in cultured cells, the MGBDs have received little attention as anti-HIV agents because many of the distamycin-related compounds are expensive and toxic to cells (39). Likewise, to our knowledge, MGBDs have not been tested for activity against simian immunodeficiency viruses or the lentiviruses that infect nonprimate mammals, though the nucleotide composition of these viral genomes are nearly identical to that of HIV-1(8). Much more emphasis has been devoted to the synthesis of distamycin conjugates and analogues as anti-HIV drugs with the goal of targeting the ligands to defined features of the viral life cycle (12). For example, several distamycin-related polyanionic conjugates are inhibitors of HIV-induced cell killing and mechanistic studies suggest that the inhibition is due to the inhibition of virus attachment to CD4+ susceptible cells (40). Another series of distamycin derivatives was designed to target the binding sites for several cellular transcription factors and a combination of these ligands inhibited HIV replication in lymphocytes (41). However, as discussed by Tutter and Jones (42), the most significant problems associated with these analogs is the ease with which HIV-1 evolves to evade chemical inhibitors and the fact that no single sequence in the HIV genome is absolutely essential for virus replication. The studies described in this report point to an alternative global specificity of the MGBDs for the HIV provirus DNA that is based on the characteristic A-bias of the lentivirus genome. The bias dictates the unusual composition of essentially all lentiviral proteins and may also serve to enhance genetic variation and curvature of the HIV genome (8,9). Consequently, the bias is considered to be critical for HIV phenotype making it unlikely that HIV variants that lacked this genome-wide sequence feature would appear during the course of an infection. Drugs whose anti-viral action is dependent on this global bias may thus provide a novel class of anti-lentiviral agents.

Nucleosome-length sequences with AT site densities as high as those seen in HIV are rare in the eukaryotic genome as implied from the results in Figs. 3, 5, and 7 which show that the MGBDs did not noticeably inhibit nucleosome assembly or promote nucleosome loss on genomic DNAs from human, chicken or yeast. The unusual nucleotide bias of HIV is the major driving force behind this specificity for it dictates the high frequencies of AT sites and their narrow minor groove character. All lentiviruses exhibit the strong A-bias and display little or no selection against the high levels of AT-trinucleotides (8) and tetranucleotides (Fig. 2) in their coding regions. In fact, the frequency of AT sites with narrow minor grooves in HIV is higher than in random sequence DNA (Fig. 2) which may be related to the propensity of the lentiviral reverse transcriptase to elongate preexisting oligo-A tracts with A at the expense of G (43,44). In contrast, less than 0.1 % of the sequences in the primate subdivision of GenBank display an A-bias that is extreme as that seen in HIV (8). In addition, oligonucleotide-length AT sequences are counter selected in cellular gene(45). Thus, the unusual viral bias unopposed by selection results in a high genomic levels of AT sites which are expected to be binding sites for the MGBDs. DNase1 footprinting studies confirmed this expectation by showing that DAPI protected nearly all of the AT sites > 2-3 bp in the HIV pol fragment 1. The compositional bias of the viral genome also favors AT sites with narrow minor grooves since homopolymeric A-tracts comprise a relatively large fraction of the AT sites. In addition, AT-containing oligonucleotide sequences with TA steps are underrepresented in HIV DNA as compared to random sequence and to AT sites without TA steps (Fig. 2). This viral characteristic is also unusual since the AT-containing oligomeric sequences in cellular DNA are enriched in TA but not AT dinucleotides (48). This feature contributes to the specificity of the ligands for the HIV genome since the narrow minor groove of AT sites is required for high affinity binding of most of the MGBDs (13, 27, 28). In addition, our previous studies have shown that extent of drug-induced inhibition of nucleosome assembly on synthetic repetitive fragments followed the order (CGGGAAAACC)n>(CGGGAATTCC)n >> (CGGGATATCC)n > (CGGGTTAACC)n ~(CGGGACAACC)n (13). This order roughly follows the relative frequencies of these AT-sequence motifs in the HIV-1 genome (Fig. 2). Thus, the A-bias favors the HIV genome that is rich in clusters of AT sites that preferentially bind to MGBDs and these features are responsible for the marked specificity of the MGBDs for the disruption of chromatin (13, 14).

The aromatic diamidine MGBDs including berenil, DAPI, and pentamidine are a group of related low molecular weight drugs that bind to AT-rich sequences (35-38). As shown in Figs. 3 and 4, DAPI was the most effective drug tested at inhibiting nucleosome formation on HIV DNA. DAPI was also more effective than distamycin and Hoechst at blocking the assembly of nucleosomes and negating the effects of intrinsic DNA curvature and anisotropic bendability in studies that used other naturally occurring and synthetic DNA sequences with multiple closely spaced A-tracts with narrow minor grooves (13,14). The high activity of DAPI as compared to distamycin and Hoechst has been attributed to the short length of this ligand and its absolute specificity for A/T sites with narrow minor grooves (13). This specificity may ensure that only particular minor grooves that give rise to curvature and anisotropic bendability are occupied by the drug. Berenil shows the same AT-specificity as seen with DAPI although the binding affinity of this ligand for AT sites is lower than for DAPI (27). This observation may have clinical implications since berenil was active in disrupting HIV-containing nucleosomes in vitro and in vivo (Figs. 3, 4, 7, 8) and this drug is currently used to treat a variety of parasitic infections in humans and other mammals (36, 46,47). In addition, in contrast to many of the MGBDs, berenil and pentamidine display low toxicity and both drugs have been reported to be non mutagenic in mammalian cells (48-50). Distamycin and Hoechst 33258 have most often served as the starting point for synthesis of most MGBD conjugates and analogs (12, 49). A similar approach could be adopted for the synthesis of conjugates of the structurally simpler drugs such as berenil, pentamidine, or DAPI in order to optimize the specificity of these compounds for the clustered AT-sites in the HIV genome. In addition, berenil has previously been used to target cytotoxic platinium to AT rich regions in the genome (51). Such conjugates targeted to the bone marrow using bone marrow targeting systems (52) might also be useful in promoting selective nucleosome disruption and lethal mutagenesis of the integrated HIV provirus.

Dimers of distamycin and neutropsin connected by a diammine bridge displayed a higher affinity for long (N=8)A/T sites in synthetic DNA than the corresponding monomers. These and similar bridged molecules show heighten anti HIV and 2 activity in culture cells relative to the monomers (53,54). However, such long A/T tracts are rare in the HIV genome. Rather, the A richness of the HIV genome is due principally to A-tracts that are 3-5 bp in length and these tracts are frequently arranged in a 10 bp periodicity, a period that is coincident with the pitch of the DNA helix (9). In addition, the specificity of the MGBDs for promoting nucleosome disruption requires long (>120 bp) stretches of DNA containing such phased A-tracts (13,14). These considerations raise the interesting possibility that dimers or higher multimers of the smaller MGBDs such as berenil with linker lengths that correspond to the helical repeat might be more effective and specific than the monomers for HIV nucleosome disruption.

The MGBDs prevented nucleosome formation on the five segments of HIV pol studied in this report and it is likely that the drugs would exert this selective inhibitory action on the bulk of HIV chromatin. It is also clear from the data in Fig. 5 that the MGBDs promoted a preferential loss of histone octamers from HIV DNA in high salt since the DNA released from these particles comigrated with free DNA on PA gels and failed to bind to nitrocellulose filters which bind protein-DNA complexes. Whether the drugs induced a loss of octamers from HIV sequences in yeast is more uncertain. The results in Figs. 7 are consistent with such a loss but the presence of loosely bound disrupted nucleosomes at these sites cannot be formally excluded. Perhaps the most direct evidence supporting nucleosome loss is provided by the results in Fig. 8 which show that the drugs reduced the number of superhelical turns of HIV-containing plasmids. However, the human SWI/SNF complex has been reported to promote a reduction in DNA supercoiling per nucleosome without causing nucleosome loss (61) so that the data in Fig. 8 do not prove that the drugs cause nucleosome loss in yeast.

Nucleosome loss and/or disruption has been correlated with cell death, promoter activation, changes in higher order chromatin structure, and enhanced nuclease accessibility (32,33). The importance of chromatin disruption in the mechanism of action of anti-cancer DNA-binding drugs has also recently been highlighted by Gurova (56). Disruption of HIV chromatin by MGBDs is expected to be detrimental to the expression of the provirus and perhaps even to the cell that contains this integrated sequence since AT sites with narrow minor grooves are a major determinant for nucleosome stability and positioning (57 58). In addition, the HIV genome is highly deficient in CpG dinucleotides and methylation of this dinucleotide enhances nucleosome stability (7,8,59,60).

The results in Figure 4 show that the MGBDs have no detectable effects on the assembly of nucleosomes on genomic DNA illustrating the specificity of the drugs for HIV sequences. However, it is likely that there are drug sensitive genomic sequences and it might be informative to understand the nature of these sequences. We previously studied this question by first performing a computer search of all sequences in the bacterial and invertebrate subdivision of GenBank in order to identify all genes that have an unusual nucleotide composition similar to HIV (61). The results revealed that about 80% of the identified genes coded for membrane protein antigens (virulence factors) from mammalian pathogens and the amino acid composition of these proteins were similar to those from HIV. We also demonstrated that the MGBDs inhibited nucleosome formation on these sequences with a drug specificity similar to that displayed by the HIV sequences analyzed in Figure 4 (62). These membrane antigen genes included those from *Pneumocystis jirovecii, Entamoeba histolytica,* and *Plasmodium vivax*. Whether this effect is involved in the anti-pathogenic action of the MGBDs in these organisms is a topic for future investigation.

Human fragile sites are segments of chromatin that fail to compact during mitosis and are seen as weak staining gaps and breaks in metaphase chromosomes (63,64). A subset of these sequences are AT rich and the expandable microsatellite FRA 16B is induced MGBD (64). This site was predicted to be packaged into MGBD sensitive nucleosomes and this prediction has been confirmed by experimental studies (14,65). Berenil has been recommended for detection of this site as compared to the other MGBDs in metaphase chromosomes because of it’s lower cost, it induces higher levels of the site and has no negative morphological effect on metaphase chromosomes in optimal concentration (66). The possibility was considered that the drugs might promote the breakage of HIV sequences in yeast as a result of the chromatin disruption effect but the data described in the Results Section suggest that this is not the case at least under the conditions used in this work. Thus, although the consequences of nucleosome disruption by the MGBDs remain to be clarified, they point to the need for future studies on the analysis of their effects on HIV infected vs non infected CD4+human cells in terms of cell viability, proviral sequence integrity and packaging of the sequences into nucleosomes. The results of such studies might then provide an assay for the optimization of desired effects using existing MGBDs, and testing new drug derivatives and drug multimers.

## ACKNOWLEDGMENTS

The author wishes to thank Irwin Tessman, P.T. Gilham and L.Weith for reviewing the manuscript and for helpful discussions during the course of these studies.

